# Feed, Flee, Or Fact-Find: Modeling Latent Cognitive-mechanistic Responses to Heterospecific Alarm Calls for Foraging Cardinals

**DOI:** 10.64898/2026.10.12.758620

**Authors:** Nich W. Martin, Po-An Chen, Scarlett Howell, Kathryn Sieving

## Abstract

Heterospecific alarm calls allow eavesdropping animals to obtain information about predators from other species, but how receivers translate different forms of alarm information into defensive behavior remains poorly understood. Signals may differ in whether they elicit rapid, relatively stereotyped responses or permit more graded evaluation of threat. We investigated this possibility by exposing foraging Northern cardinals (*Cardinalis cardinalis*) to playbacks of tufted titmouse (*Baeolophus bicolor*) calls associated with different predator contexts, including graded and immediate threats. To characterize responses, we developed a Bayesian latent-risk model that integrated three behaviors: feeding, vigilant looks, and departure, to estimate a shared dimension of perceived predation risk. Inferred risk increased from near-baseline levels under contact calls to intermediate levels under owl-associated call and the highest levels under ZZ calls. Increasing latent risk was associated with reduced expected seed intake and higher probability of complete feeding suppression. Vigilance showed a nonlinear relationship with inferred risk, increasing at intermediate levels before declining under the highest-risk treatment in favor of departure. This pattern is consistent with the possibility that receivers continue gathering information under intermediate threats but reduce vigilance when responding to more urgent danger. Our results show how heterospecific alarm calls map onto distinct, coordinated behavioral responses. Although not established here, we suspect that differences arise because of selection for rapid canalized responses to urgent threats and for more flexible cognitive evaluations of graded threats.

## INTRODUCTION

Animal foraging decisions involve competing demands. The balance between food acquisition and predator avoidance can depend on the energetic value of available resources, the costs of foraging, and the likelihood and consequences of encountering a predator (Lima and Dill, 1990; Brown and Kolter, 2004). Because these factors vary across time and different environments, animals must adjust their behavior to changing conditions (Lea et al., 2020). Such adjustments may include changes in feeding rate, vigilance, and the decision to abandon a foraging site (Brown, 1999). Importantly, these behaviors do not necessarily respond to predation risk in parallel (Lima and Bednekoff, 1999). For example, vigilance may increase when a forager detects a potential threat but may also decline when other defensive responses become more advantageous (Lima, 1987). Moreover, behaviors can be largely state-dependent (McNamara and Houston, 1987). For example, an energetically depleted individual may remain in a relatively poor food patch rather than explore alternative patches when the energetic costs of travel outweigh the expected benefits of discovering a higher-quality patch; whereas a well-fed individual can invest in visiting multiple patches such that they can then be exploited in turn according to their marginal value (Olsson, 2001). However, foragers rarely have perfect information (Stephens and Krebs, 1986); thus, understanding how animals balance feeding and predator avoidance requires considering not only the true risks they face, but also the information available to them when making behavioral decisions.

Animals can acquire information about predation risk through direct observation and through the behavior and vocalizations of other individuals (Jones et al., 2024). In particular, eavesdropping on heterospecific alarm calls allows animals to obtain information about potential predators without direct observation (Magrath et al., 2015). Such social information can elicit immediate anti-predator responses, influence foraging and habitat use, and provide opportunities to learn about predators (Hetrick and Sieving, 2012; Magrath et al., 2015; Huang et al., 2012; Jones and Sieving, 2019). However, responding to an alarm call also entails a decision: fleeing in response to an unreliable or non-urgent signal can impose unnecessary energetic costs, whereas failing to respond to reliable warnings may increase the probability of predation. Therefore, receivers must use social information in ways appropriate to the threat being communicated and behave in ways that maximize fitness and survival.

While models of information use can illuminate what choices animals make and why they shift between risk-prone and risk-averse options, they largely treat signals as single, static events (Ravignani and de Reus, 2019; Sayfarth et al., 2010). Incorporating the temporal architecture of cues is critical, as it directly influences the phenology of risk-perception updating and subsequent behavioral responses. Animals produce and utilize instantaneous, referential signals—distinct acoustic categories for distinct contexts (e.g., a hawk flight alarm)—as well as sequential, graded signals that rely on variable numbers of repeated acoustic elements to encode threat urgency (Sieving et al. 2010). Whereas instantaneous signals can be decoded rapidly, sequential signals require a temporal window to "unwind" their full repetition of elements to communicate urgency, or proportional processing, governed by Weber’s law (Akre & Johnsen 2014).

Kalkman (2017) provides a rigorous evolutionary framework for this acoustic dichotomy, explaining how signal structure determines the causal-explanatory pathways engaged by the receiver. For instantaneous, high-urgency alarms, natural selection installs phylogenetically canalized, reflex-like responses at the population level, where the physical signal acts as an immediate triggering cause that bypasses slow, decoupled cognitive processing to prompt rapid escape. In contrast, the processing of sequential, graded alarms relies on ontogenetic cognitive flexibility at the individual level. Here, the cumulative associated information of the signal serves as a structuring cause, allowing the receiver to form a decoupled mental representation— buffering the incoming social cue to actively weigh it against internal hunger and personal scanning before executing a flexible, context-dependent response. Consideration of the *kind* of signal (instantaneous vs. sequential) and its fitness value shapes understanding of the processing pathways engaged (autonomic vs. cognitive), and ultimately the risk-appropriate response.

One of the challenges for understanding whether receivers treat signals as high-vs. low-urgency calls, is that perceived predation risk is not directly observable (Crane et al., 2024). Instead, researchers must infer risk from behavioral responses that may provide incomplete or ambiguous evidence about an animal’s assessment of its environment. Changes in feeding, vigilance, and departure from a foraging site are all potential responses to danger, but they do not necessarily change in the same direction or at the same rate as perceived risk increases (Jones et al., 2024). Consequently, treating any single behavior as a direct measure of perceived risk may obscure important differences in how animals respond to alarm signals.

Multivariate latent approaches offer a way to address this challenge by integrating multiple behaviors to estimate a shared, unobserved dimension (Muthén, 2002), specifically of perceived risk. Rather than assuming feeding, vigilance, and fleeing are interchangeable measures of threat, these types of models allow each behavior to exhibit a distinct relationship with latent risk, providing a framework for comparing alarm signals along a common latent dimension, while accounting for differences in the behavioral expression of a threat. This approach links the study of signal architecture to the broader question of how receivers translate information about predation risk into coordinated behavioral responses.

Here, we conducted a rigorous field experiment capitalizing on the well-known anti-predator communication system of birds in family Paridae. Using situationally specific anti-predator calls of a vocally complex community informant species, the tufted titmouse (*Baeolophus bicolor*), and broadcasted those calls to a highly competent eavesdropper, the Northern cardinal (*Cardinalis cardinalis*), we document decision-making response types and their phenology in relation to latent risk-perception in a foraging context. Using three distinct behaviors (feeding, vigilance, and departure time) we developed a Bayesian latent-risk model that integrated those behaviors to estimate the relative perceived risk across treatments.

We expected call treatments to differ in their inferred risk. Based on results from state-dependent optimal behavioral model (Supplementary Materials), we expected treatments associated with more dangerous predation contexts corresponding to stronger suppression of feeding and greater likelihood of departure. Similarly, we expected vigilance to exhibit a non-monotonic relationship with the inferred risk, increasing under intermediate threat conditions, but declining under the most urgent threats. By integrating our generative model with multiple behaviors and a latent-risk framework, we sought to characterize how different alarm-call treatments correspond to perceived risk and to examine how the behavioral expression of that risk varies across heterospecific calls.

## METHODS

### A model communication system

Avian anti-predator calls are ubiquitous in nature, and they can generate taxonomically wide-reaching responses in diverse eavesdropping communities. The complex anti-predator signaling system in Family Paridae is among the best studied of these communication systems (Hetrick & Sieving 2012; Krams et al., 2012). Parids live year-round in small family groups but are also highly heterospecifically social. They lead winter foraging flocks (Farley et al. 2008; Contreras & Sieving 2011) and can generate large and diverse mobbing aggregations around discovered predators via vigorous mobbing vocalizations (Sieving et al. 2006; Sieving et al. in press). As such, parids influence a diverse set of eavesdroppers in local communities they inhabit. Parids have two major call types they use to identify immediate and future predation risk. The ZZ (seet, or high see) call is a high-pitched series of tones (8-9 kHz) typically given when an actively hunting or attacking hawk is spotted near a parid family group or mixed foraging flock (Pieplow 2017). The call causes nearby small birds to rapidly escape to cover (ditch to ground or flee the area) or to freeze in place to avoid being attacked (Jones & Sieving 2019). As such, ZZ calls are likely classified as instantaneous signals. Additionally, most parids produce ‘chick-a-dee’ (or CD) calls taking the form of several modulated tones (chick notes in Sieving et al. 2010) preceding a string of high entropy D notes that varies in number (Figure 1; Hetrick & Sieving 2012). Long CD calls with many D notes are produced when parids encounter a stationary predator (small owl or hawk) perched or loafing in their vicinity. The number of D notes increases with the dangerousness of the species of avian predator. Small owls that consume many small birds generate longer D note sequences per CD call than large, clumsy owls that rarely prey on small birds (Templeton & Greene 2005). In this way, the CD calls convey the magnitude of danger represented by the encountered species via gradations of D note number. Given that detecting differences in urgency between CD calls is governed by Weber’s law, or a receiver’s sensitivity to changes in urgency that are encoded within graded signals, this call type likely functions as a sequential signal. Cognitive involvement is essentially required to count D notes to accurately gauge threat perception (Nieder 2020).

**Figure 1.**
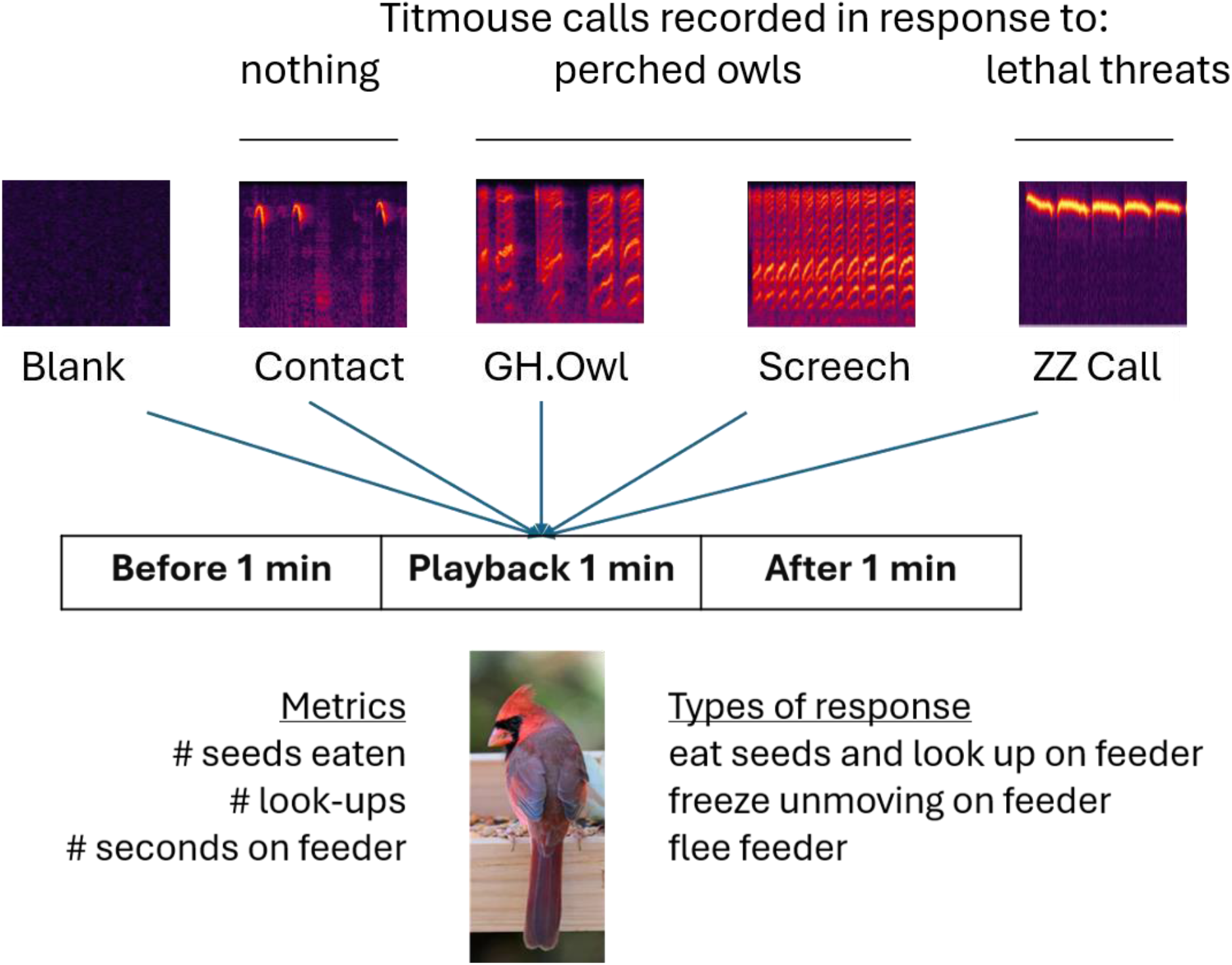
The diagram summarizes the experimental design and acoustic architecture of heterospecific playback treatments. One of 5 playback treatments was played for one minute during minute 2 of 3 observation minutes from a speaker at 3 m height above a platform feeder. Behavioral metrics (counts) recorded for a lone Northern cardinal in all cases included seeds eaten, head lookups, and time-to-flee. Categories of behavior noted included active feeding and looking, freezing on feeder, and fleeing from the feeder. Sonograms at the top depict approximately two seconds of each vocalization type recorded previously during live predator presentations to titmice (blank (silence), contact calls, chick-a-dee calls (CD) given to a perched Great Horned Owl, chick-a-dee calls given to a perched Eastern Screech Owl, and calls with repeated Z notes (ZZ call) given by titmice to live predators up close representing potential immediate lethal threats to the caller(s). Frequencies in sonagrams range from 1 to 9 kHz.

**Figure 2.**
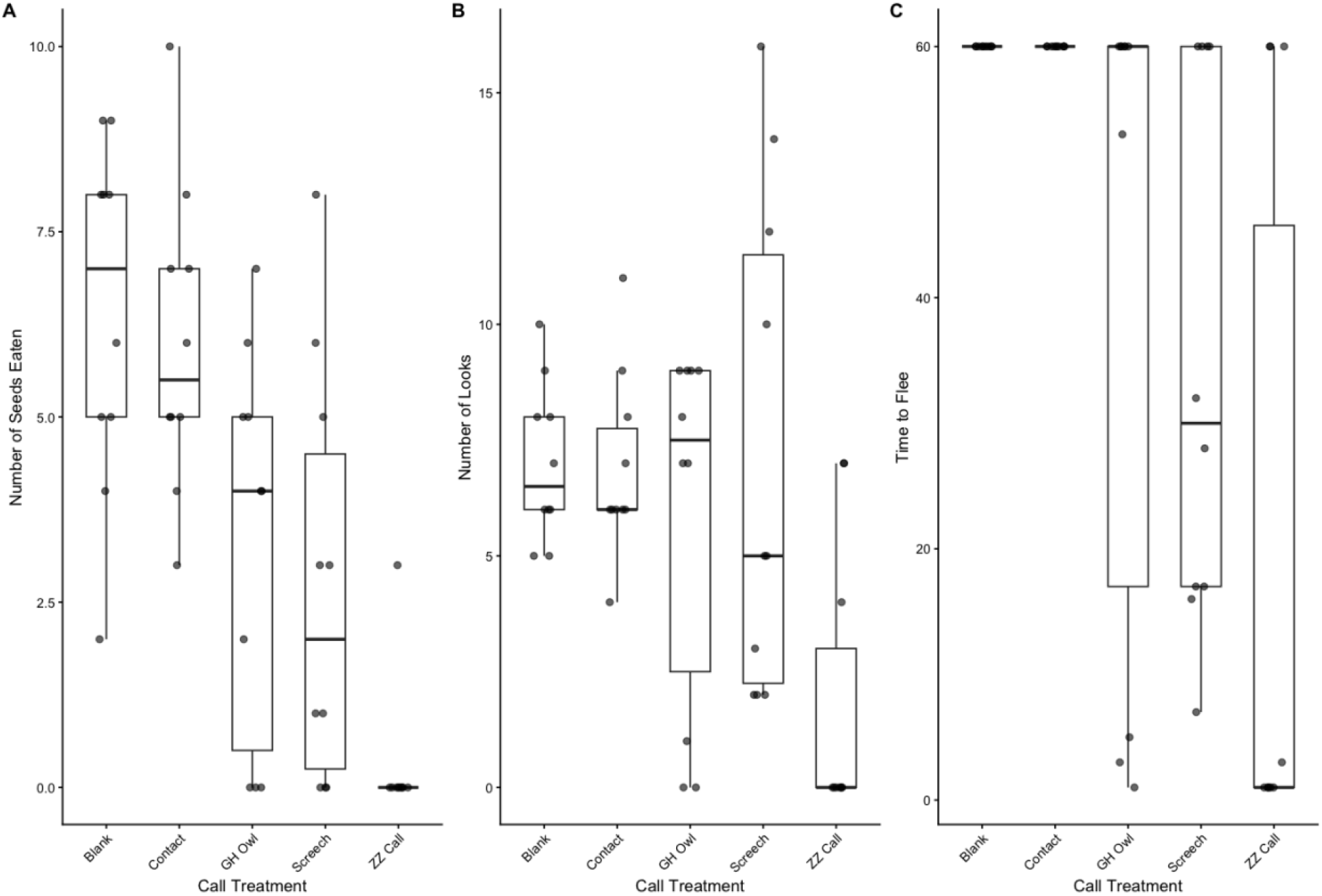
Box plot and points of raw data for number seeds eaten during callback period (A), number of looks taken (B), and time the bird fled the feeder (C). Birds that did not flee during the trial were censored at 60 minutes.

### Field experimental design

We designed a field experiment using playbacks of ZZ alarm calls, high risk CD (many D notes), and low-risk CD (short D sequences) calls to lone Northern cardinals foraging on platform feeders (**Figure 1**). We established 10 independent platform feeders in suburban yards in Gainesville FL at least 500m (and up to several km apart) and kept them filled with fresh *ad libitum* black oil sunflower seed over 3 months in 2006. Playback stimuli were constructed using calls taken from recordings made during two experiments and in the field. In the two previous experiments, live predators were either presented to 3 free-flying titmice in an aviary (Hetrick & Sieving 2012) or to a single live titmouse confined to a small cage (Sieving et al. 2010). In the former, predators were tethered to a perch, and 5 different flocks of 3 titmice were exposed. In the latter, live predators were presented while confined in a plexiglass box at 1 meter from the confined bird (N = 15 titmice). In both, the first 2 minutes of recordings of titmouse vocalizations following an unveiling of the predator (remotely lifting a pillowcase covering; see Hetrick 2006 for details) were used as source files for this study. Each recording constructed for this experiment (using Avisoft SASLab Pro) presented a series of calls taken from files associated with the appropriate stimulus. Chip notes, and all CD calls (to Screech and GH Owls) were taken from recordings made in Hetrick & Sieving (2012). Several ZZ call sequences were taken from birds responding to a large housecat presented at close range in Sieving et al. (2010), and two recordings of repeated ZZ calls were taken from field recordings of flocks as an *Accipiter spp.* attacked the flocks (Sieving unpublished data). Each recording was standardized to have the same number of total calls, and within each treatment type we attempted to equalize the total number of notes per recording. In addition to Contact call (soft chip notes that titmice produce when not agitated), 2 CD call and 1 ZZ call treatment, we used a procedural control (silent recording), making 5 total playback treatments **Table 1**). Five variants of each playback type were made, that mixed and matched calls taken from different source files, such that each variant was unique (in terms of the individual birds represented) but had the same structure (calls and/or note numbers). Each feeder received a randomly selected variant for each treatment type (without replacement). In preliminary analyses, there was no detectable signature of variant ID, and this factor was dropped in final analyses. Each feeder received each of the 5 treatments on different days at least 5 days apart in a Latin squares design that eliminates order effects in analysis (**Figure 1**; **Table 1**).

**Table 1.** List of playback treatments, their description, signal type, and predicted threat level. See sonagrams in. **Figure 1**.

| Treatment | Description | Signal type | Threat level & ecology |
|---|---|---|---|
| **Blank** | Blank recording playing from a speaker | Procedural control | Baseline environment with no alarm stimuli |
| **Contact** | Soft contact calls naturally produced by titmice when not agitated | Treatment control | Negligible threat; indicative of normal, safe social foraging |
| **GH Owl** | Low-intensity chick-a-dee (CD) calls given to a live, perched great horned owl | Graded with fewer D-note repetitions / CD call than Screech | Intermediate-low threat; great horned owls are large predators that rarely target small birds |
| **Screech** | High-intensity CD calls given to a live perched Eastern screech owl | Graded (structuring) with a higher count of D-notes / CD call than GH Owl | Intermediate-high threat; screech owls are small, lethal predators of small songbirds |
| **ZZ Call** | High-pitched series of tonal Z notes given to attacking hawks in flight or a cat at close range | Instantaneous (triggering) comprised of 3-6 repeated tones (8-9 kHz) | Highest threat; signals imminent mortal danger |

### Protocol

An observer (Scarlett Howell in all cases) set up a battery boosted speaker on a painter’s pole raised to 3m and leaned against a mature oak tree near the feeder. All feeders had similar placement on suburban lawns with at least one shrub and a mature oak tree near the feeder (< 10m away). Speakers were connected to an iPod Shuffle (first generation) with audio cords and the observer sat in a small chair 20 meters away from the feeder with 10 x 50 power binoculars. Northern cardinals generally stay on feeders for one to several minutes at a time eating successive seeds and frequently looking around for predators while cracking and consuming each seed. The observer waited at least 5 minutes and sometimes longer for a cardinal to settle on the feeder and begin eating. Once settled, the observer cued a pre-selected treatment playback variant and started a timer. For one minute she counted seeds eaten and look-ups, then initiated the playback and continued counting for one minute. Then after one minute of stimulus playback, the observer continued counting for one final minute. If a cardinal left the feeder during one of the periods, then the number of seconds on the feeder in that period was noted.

### Behavioral response metrics

Cardinals tend to sit in one place on a platform feeder, eating one seed after another by dipping the head to pick one up and then raising the head (look-up) while manipulating the seed to extract the meat. During three periods of observation, before, during and after exposure to playback (**Figure 1**), we recorded counts of whole seeds eaten, head raises (look-ups), and time spent on the feeder. We also recorded if a bird left the feeder and after how many seconds during the trial minute that happened. All birds stayed put on the feeder eating seeds for the full pre-trial minute (60 seconds), providing a control (before) for seeds eaten and look-ups per minute for each bird for comparison to the trial (playback) minute. Some birds stayed on the feeder for all three minutes in all treatments, but if a bird left a feeder, it was during the trial minute (except for 1 individual who left in period 3; see Sieving et al. 2026). No birds returned to the feeder during observations if they left.

## ANAYSIS

First, we developed a generative model using a combination of dynamic optimization (Clark and Mangel, 2008) and forward simulation to determine the qualitative changes in the total number of seeds eaten, looks taken, and time to flee for increasing levels in the rate at which risk increases through time. This state-dependent model assumes fitness increases with the total number of seeds eaten over the trial period and decreases as the immediate perception of risk increases. Mathematical details of the model can be found in our supplementary materials.

We parameterized a Bayesian latent risk model to treat the underlying threat types and urgencies as an unobserved (latent) variable. The observed behaviors are modeled as joint outputs caused by that latent risk. Together these approaches provide a rich, probabilistic interpretation of how varying architecture of the 4 titmouse call types translate into incremental risk perception and behavioral adaptation (compared to a control).

### Latent risk model parameterization

We determined relative perceived risk of each call type *k* using a Bayesian latent variable model with multiple behavioral indicators and hierarchal effects. Behaviors were linked through a shared predation-risk axis, and each call type was assigned a latent perceived-risk parameter *r_k_* anchored to the Blank playback:

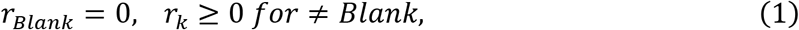

with higher *r_k_* representing greater perceived risk. Based on the results of our generative model, we assumed the number of seeds eaten *SE_n_* and time-to-flee *T_n_* would decrease with increasing risk, while the probability of fleeing *F_n_* would increase. We also assumed the number of looks would increase at low-intermediate risk levels but decrease at some threshold as birds begin to leave the feeder earlier than they would at lower levels of perceived risk, minimizing the available time to look. We also assumed a state-dependent trade-off between the number of seeds already eaten and perceived risk of predation. Thus, we included number of seeds eaten prior to playback *PSE_n_* as a predictor of fleeing behavior. For better sampling, we chose to center *PSE_n_* such that

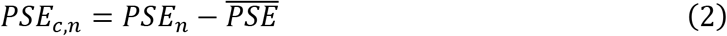

Because each feeder was at a different location and may differ in background risk and foraging conditions, we included feeder-specific varying intercept for all behaviors such that,

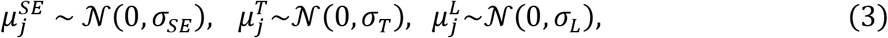

where *j* ∈ {1, …, *J*} represents feeder index. These terms shift the mean feeding rate, baseline tendency to flee, and mean look rate, respectively.

### Fleeing

We modeled leaving as a discrete-time hazard, censored at 60 seconds, such that for trial *n* at second *t*,

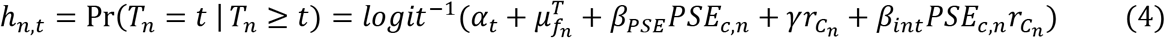

where *α_t_* is a time-specific baseline logit-hazard, *f_n_* is feeder identity, and *C_n_* is call type. The interaction coefficient *β_int_* predicts how baseline feeding changes fleeing likelihood as perceived risk increases.

### Feeding

Number of seeds eaten during playback was evaluated using a negative binomial likelihood such that,

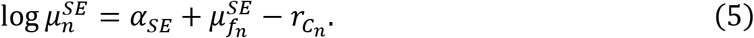

We also included a zero-inflation term to allow “no feeding” trials beyond what the negative binomial would predict to account for “freezing” states in which a bird would not feed but also remain on the feeder:

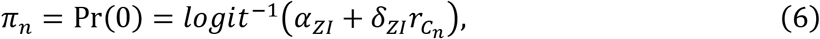

thus, allowing risk to reduce feeding by lowering the expected seed count and increasing the probability of complete feeding cessation.

### Lookups: exposure-adjusted peaked risk response

Lookups were also modeled using a negative binomial distribution, but because the number of looks from our dynamic optimization model were predicted to be non-monotonic with respect to risk, we used a quadratic term with an exposure offset for the time on feeder:

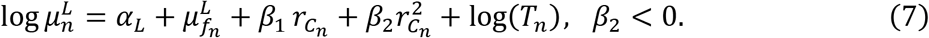

The model was fitted in Stan (release 2.39.0) using Hamilton Monte Carlo with weakly informative priors:

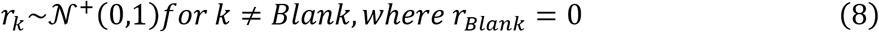

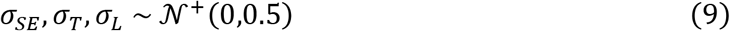

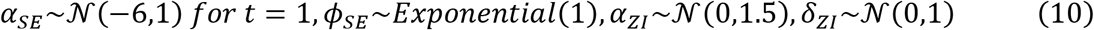

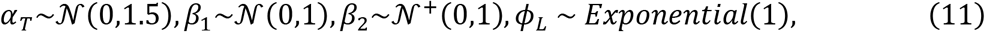

where *φ_B_*is the error term of the negative binomial likelihood for response *B*, where *B* ∈ {*SE*, *T*, *L*}.

## RESULTS

The latent risk posterior (anchored at Blank = 0) showed a clear and interpretable gradient. Contact was only slightly elevated relative to Blank, GH Owl was intermediate, Screech was higher than GH Owl, while ZZ call was substantially higher than all call types (Table 2; Figure 3). This ordering is consistent with the expectation that Contact calls do not signal predation risk, GH Owls are relatively low risk predators compared to Screech owls, and the ZZ call represents the highest risk, representing clear and immediate danger from a nearby mortal enemy (Sieving, et al. 2010; Hetrick & Sieving 2012).

**Figure 3.**
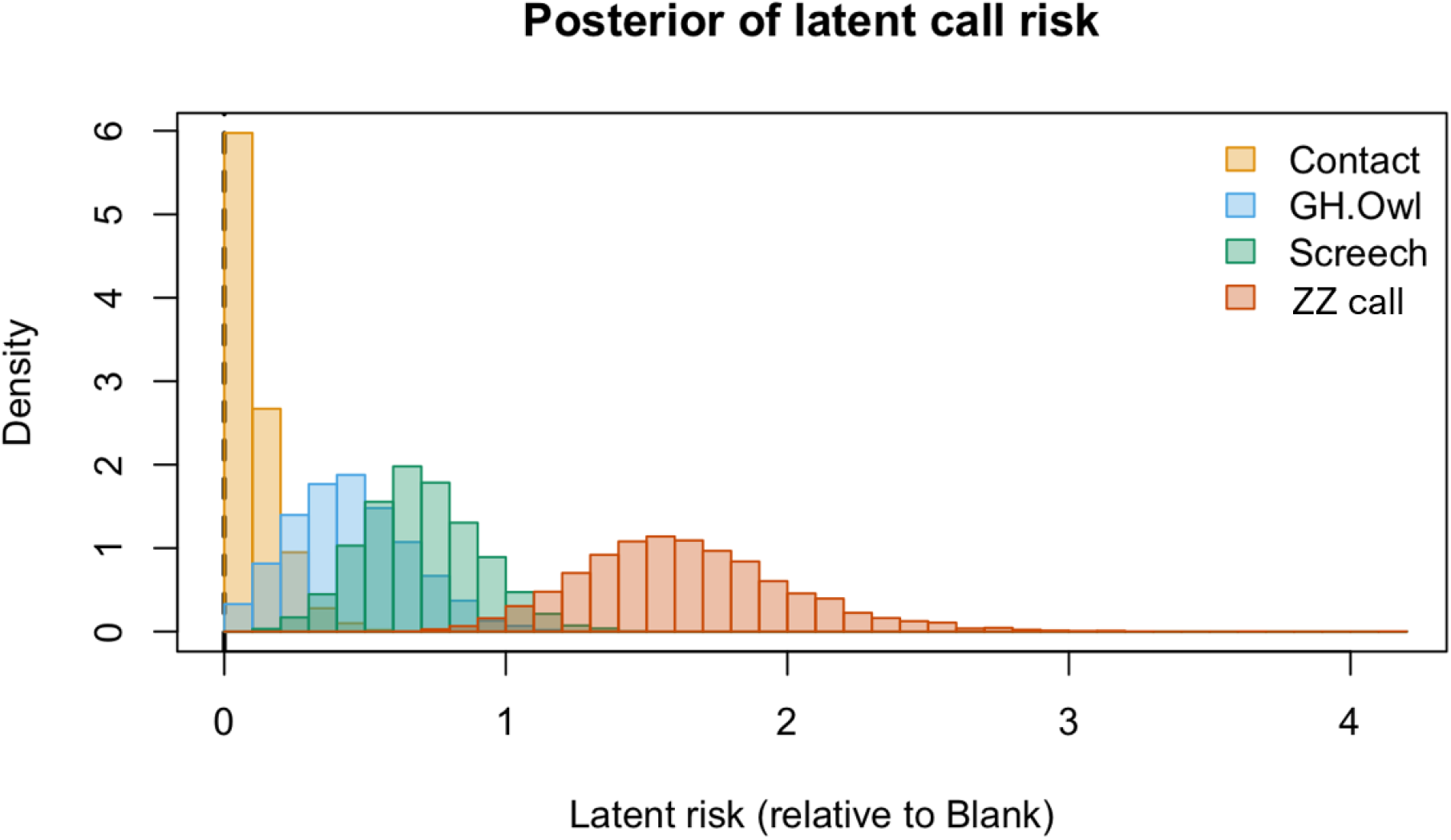
Posterior distributions of latent perceived risk (*r_k_*) for five parid call types. This plot shows the clear and interpretable risk gradient inferred from the Bayesian latent variable model, with the Blank playback anchored at zero (dashed vertical line). Latent risk estimates were derived from three behavioral axes: feeding suppression, vigilant looks, and fleeing hazard. The results show a progression of perceived risk from Contact calls (near zero) to intermediate-risk owl CD calls (Great Horned Owl < Screech Owl) and finally the highest-risk (ZZ) calls.

**Table 2:** Bayesian median risk value estimates (r =) from Bayesian model (± 90% credible intervals; 5-95%) for each treatment: Latent posterior, feeding suppression, expected seeds eaten and fleeing (see text for lookup results).

| **Behavior** | **Blank** | **Contact** | **GH.Owl** | **Screech** | **ZZ call** |
|---|---|---|---|---|---|
| **Latent risk posterior** | 0 (anchor) | 0.079 (0.006-0.285) | 0.435 (0.130-0.823) | 0.689 (0.375-1.054) | 1.609 (1.090-2.319) |
| **Feeding suppression** | 0.103 (0.038-0.209) | 0.119 (0.044-0.231) | 0.190 (0.078-0.342) | 0.255 (0.104-0.432) | 0.591 (0.198-0.853) |
| **Expected seeds eaten** | 5.764 (4.432-7.598) | 5.116 (3.826-6.751) | 3.341 (2.129-4.883) | 2.381 (1.505-3.602) | 0.515 (0.148-1.213) |
| **Fleeing** | na | na | 0.343 (0.237-0.505) | 0.437 (0.307-0.606) | 0.82 (0.585-0.953) |

### Feeding suppression

The zero-inflated feeding sub model detected strong risk-dependent increases in the probability of a non-feeding bird (i.e., early Fleers and Freezers). The posterior median zero-seed probability increased along the same call-risk gradient (Table 2): Blank 0.103, Contact 0.119, GH Owl 0.190, Screech 0.255, ZZ Call 0.591. Likewise, the zero-inflation-adjusted expected number of seeds eaten declined strongly across calls (Table 2): Blank median 5.764, Contact 5.116, GH Owl 3.341, Screech 2.381, and ZZ call 0.515. Thus, the ZZ call was associated with both substantially reduced feeding and a much higher probability of complete feeding suppression.

### Fleeing

Median probability for fleeing during the trial was highest for birds receiving the ZZ call (Table 2) and was notably lower for GH Owl and Screech; 0.343, 0.437, respectively. For Blank, Contact, GH Owl, and Screech calls, the effective slope for the number of seeds eaten prior to the playback (pre-trial period) was typically negative (mean slopes: -0.169, -0.157, -0.116, - 0.090; with *P*(*slope* < 0) = 0.77, 0.77, 0.75, 0.72, respectively), suggesting higher feeding prior to playback treatment tended to predict lower flee hazard (greater persistence on the feeder). This was contrary to our prediction that state-dependent trade-offs would show a higher flee hazard, where birds who consumed more seeds prior to playback treatments would be more satiated and willing to leave the feeder. In contrast to the other calls, the effective slope for ZZ call was centered near 0, showing a high level of model uncertainty (median 0.015, -0.368-0.445), consistent with weak or possibly absent state dependence for leaving under the highest-risk warning call.

### Looking

The vigilance model implied an inverted-U relationship between latent risk and look rate (Figure 4). The posterior distribution of the turning point (i.e., peak looks due to risk) was *r*^∗^median 0.740 (5-95%: 0.271-1.315), and call types differed in whether they were predicted to fall above or below this turning point. The probability *r_k_* > *r*^∗^ was 0.022 for Blank, 0.032 for Contact, 0.189 for GH Owl, 0.439 for Screech, and 0.974 for ZZ call. Thus, ZZ call almost always occurred on the decreasing side of the vigilance curve, consistent with high attack-risk conditions eliciting reduced scanning either through fleeing or immobility.

**Figure 4.**
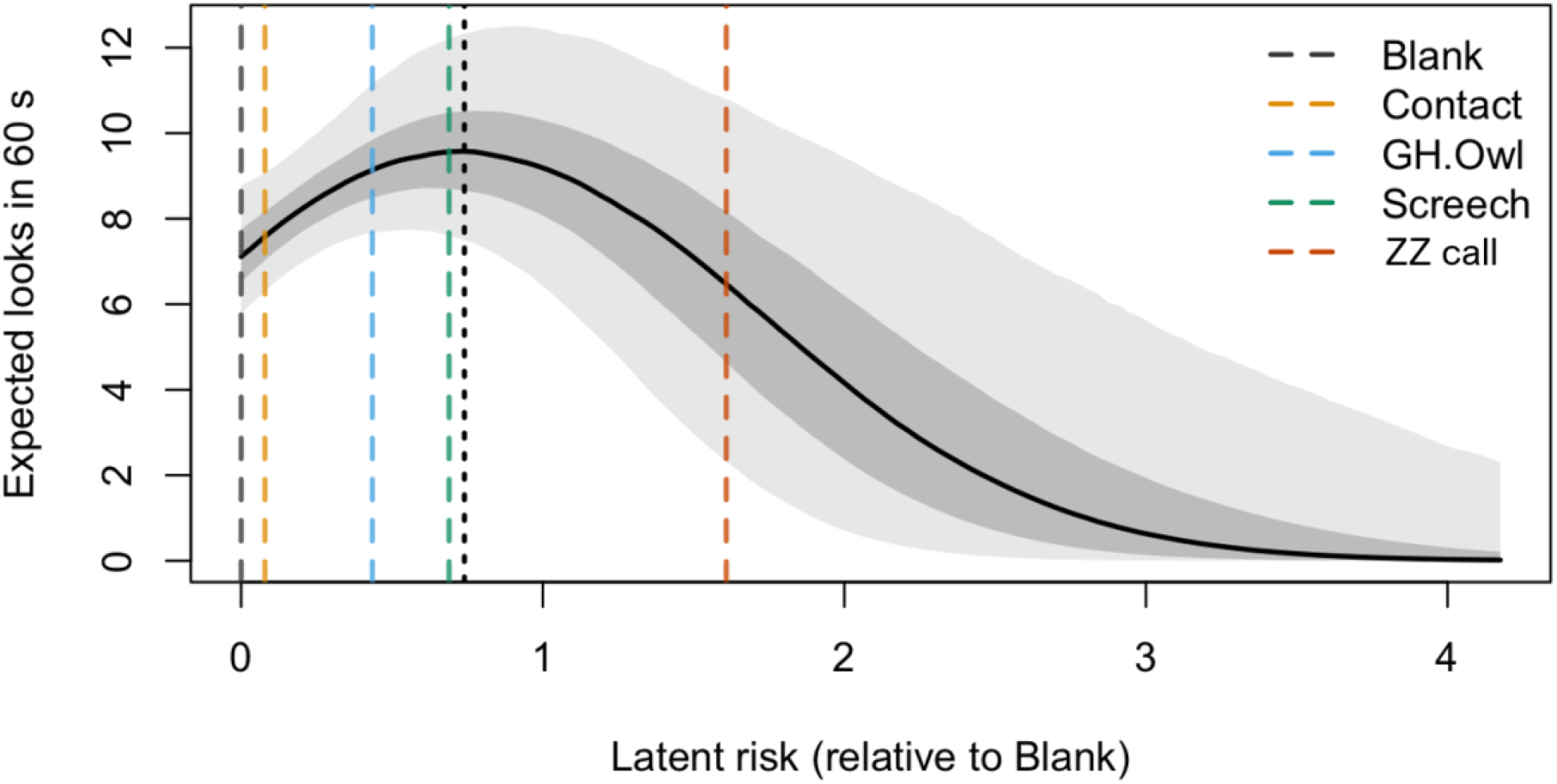
The "Vigilance Curve": The non-monotonic relationship between perceived risk and lookup rates. The model illustrates an inverted-U relationship between the latent risk of a signal and the rate of vigilant head-looks. Vigilance increases from low to intermediate risk levels as birds scan for the potential threat but decreases at high risk levels (ZZ) as birds transition into a "defense mode" characterized by either rapid departure or total immobility. The quadratic term (*β*_2_ < 0) in the negative binomial model captures this exposure-adjusted peaked response. Dark grey shading represents the 95% CI for the predicted posterior mean number of lookups against latent risk; light grey is the additive distribution of the negative binomial dispersion parameter (*φ_L_*).

## DISCUSSION

### Multidimensional behavioral responses to heterospecific alarms

Rather than responding to heterospecific alarm calls as undifferentiated disturbance signals, cardinals showed clear evidence of extracting associated, predator-specific threat information from the vocalizations of titmice (Huang et al., 2012), translating these signals into multi-axial behavioral changes. Single latent risk estimates were inferred from three behavioral axes (feeding, vigilant looks, and fleeing) and median posteriors of the latent risk for each call type showed a clear gradient ordered according to the degree of risk associated with each predator/call type—low (Contact) to increasing intermediates (GH Owl and Screech) to high (ZZ call). Responses to increasingly riskier signals were not simply an increased tendency to flee but were also expressed as coordinated distinct shifts in foraging and vigilance behavior. Thus, three behavioral axes converge to map a singular underlying risk spectrum. This demonstrates a qualitative shift in avian processing strategies across both graded threats and referential situations shifting from intermediate potential threats and immediate mortal danger.

The non-linear relationship between look-up vigilance and estimated threat level is highly consistent with the distinction between “phylogenetic adaptation” (genetically canalized, population-level reflexes) and ontogenetic flexibility (real-time, individual-level cognitive processing) in cognitive ecology (Kalkman 2017). When parsing graded, sequential alarms (such as the titmouse screech owl CD calls), eavesdropping cardinals did not exhibit a hard-wired, obligatory response. Instead, they utilized what are known as decoupled mental representations (Kalkman 2017)—representing the statistical signal-predator correlation internally so that it can be actively weighed against personal visual information (lookups) and internal energetic state. This cognitive buffer explains why lookups peak under intermediate threats: the signal serves as a structuring cause that guides real-time decision-making and prompts personal verification. In contrast, the highly urgent, instantaneous ZZ call represents a threshold transition where the physical signal acts as an uncompromised triggering cause. Under this imminent mortal danger, the prohibitively high survival cost of personal verification collapses the decoupled processing phase entirely, potentially shifting the receiver’s nervous system directly into autonomic, state-independent flight or sympathetic dominance. Interestingly, cardinals remaining on feeders during playback of Screech greatly increased scanning (and accrued seed deficit), but not for the GH owl, favoring a smaller seed deficit for this treatment. In both cases, a perched owl is not an imminent mortal threat, but the potential that a more dangerous screech owl might be watching forces a direct trade-off between attempts to locate the owl (lookups) and feeding that does not occur with the horned owl who is unlikely to attack even if it has eyes on the prey.

These findings also highlight the interplay between prey reliance on (and valuation of) both social and personal information collection (Gil et al., 2017) when dealing with graded and referential threats. Under immediate, high-urgency threats (such as a ZZ call signifying an attack in progress), the high cost of taking any time for personal verification causes a complete shift in valuation to the socially derived cue, bypassing personal scanning to initiate rapid escape or freezing. Most notably, the zero-inflated feeding submodel detected that under ZZ playbacks, feeding was sharply suppressed independent of immediate departure. This suggests that high-risk alarm information triggers a state of sympathetic dominance, where attention and motor output are physiologically redirected away from food handling, even in individuals that choose to remain on the feeder. This cognitive and behavioral reorganization occurs independently of a physically overt, classic ’freezing’ response.

### Cognitive engagement & information valuation

Here we detected how birds utilize both graded and referential signals and both personal and social information concerning predation risks. Three classes of calls, increasing in threat level, were presented – contact chip notes, CD (two gradations) and ZZ calls. The treatment associated with the highest seed intake (and near zero latent predation risk; Figure 3) was the TUTI contact call stimulus. In this case, the information available to cardinals suggested (a) that a competent predator warning system was present and (b) not being triggered by any threats. Contact calls are trustworthy ‘all clear’ signals (Sieving et al. 2010; Goodale et al. 2020) allowing birds to forage confidently with little need for additional information. Determining perceived risk under the two CD call treatments, however, is a much more difficult endeavor. As graded, sequential signals given to perched owls and climbing snakes (Soard & Richardson 2009; Suzuki 2014). For the cardinal caught in a trade-off between reducing starvation risk by continuing to feed and correctly deciding when and if to escape to survive, the individual must listen while the call sequence unwinds over time and across repeated emissions to evaluate the sequence. A forager’s sensory discrimination determines how quickly and accurately the bird can update its informed perception of risk over some number of seconds as the playback unwinds over 60 seconds in our study. This cognitive delay does, however, provide a temporal window that cardinals can utilize to collect personal information (via lookups) to verify whether a predator is present nearby (which was not in this study). Especially under the effects of the Screech, but not the GH Owl threat (Figure 4), cardinals amplified their lookup rates. Counting lookups gave us an indicator of personal information available to cardinals, just as seed counts provide a relative measure of satiety. Each of these resources (food and information) modifies perceived risks of starvation and mortality, respectively (Lima 1987; Lima & Bednekoff 1999), but because the two risks are linked, the balance of food and information intake can counter-act each other in decision-making (Bobb et al. 2025). More food should make birds more predation-risk averse, but more vigilance can reduce the perception of predation risk, and this may explain the counter-intuitive finding in the Bayesian model. It is also clear that when a trustworthy social signal indicates an immediate mortal threat, as does the ZZ call to a lone foraging cardinal, food and information gathering both become too costly to seek and the decision-making collapses to a seemingly autonomic (non-cognitive) response designed to avoid a mortal attack (flee or freeze). Our findings potentially highlight the utility of the phylogenetic (population-level autonomic) versus ontogenetic (individual-level cognitive) dichotomy when modeling prey behavioral tradeoffs in complex conditions (Kalkman 2017; Roelofs & Dayan 2022).

### Optimal foraging, state-dependence, and the autonomic shift

Field tests of optimal foraging theory consistently confirm that every foraging decision is made in the context of risks (Fletcher & Sieving 2010; Houston & Rosenström 2024). For small birds especially who are unable to carry much fat while maintaining efficient flight, starvation risk is ever present (Broggi & Nillson 2023). But so also is the risk of predation from strategically located carnivorous insects (mantids; Nyffeler et al. 2017), cryptic snakes on the ground and climbing through the canopy (DeGregorio et al. 2014), to terrifying accipiters and falcons who strike with surprise and chase them on the wing at high speeds, often from above (Beauchamp 2023). Increasing food intake lessens starvation risk perception and increasing information intake can lessen predation risk perception (Bednekoff & Houston 1994), and in cardinals as in many foragers, the two activities are in constant tension. With a full belly, birds generally become predation risk averse and vice versa (Bonter et al. 2013). Yet with more information indicating safety, they will eat more in a risky situation (Bell et al. 2009), as we observed here.

While standard optimal foraging models (e.g., Clark and Mangel, 2008) predict that highly satiated birds should flee sooner, our fleeing model revealed the opposite: increased baseline feeding intensity prior to playback (PSE) was associated with lower flee hazard and longer persistence. Rather than violating state-dependent theory, this finding likely uncovers hunger as a powerful latent state; individuals arriving highly motivated to feed tolerated significantly more risk before departing. Under the immediate threat of the ZZ call, however, this integrative, state-dependent decision rule was markedly attenuated, with the PSE slope centering near zero. This suggests a threshold-like switch where the cognitive weighing of internal hunger is completely overridden by an autonomic defense mode. This maps well onto the cognitive to autonomic threshold-like switch to an all-defensive mode where cognitive weighing of hunger is completely overridden (Kalkman 2017). Yet our results suggest some small adjustments to the exact manifestation of the autonomic response are still made based on recent information (lookups during the pretrial period) and externally cued hunger state (time of day).

### Community informants and information cascades through competent eavesdroppers

Ultimately our findings convey how information about predator identity and immediacy can be transmitted across species within a community, and that heterospecific eavesdroppers can process and integrate call differences, among referential categories and across graded scales (Sieving et al. 2010), into their own decisions regarding foraging-predation tradeoffs. Northern cardinals are demonstrably competent eavesdroppers on TUTI signals in this system (Grade & Sieving 2016) as are many other species of (especially) foliage and bark gleaning understory birds (Huang et al. 2012; Jones & Sieving 2019). It is thought that the exceptional degree of vocal diversity and versatility (Wirthlin et al. 2019) encoded in TUTI (and other parid calls; Hailman 1986; Freeberg et al. 2012) arose owing to a combination of the extended parental care characteristic of birds in family Paridae, demanding competent family communication under kin selection theory and their strong associations and collaborative problem solving with individuals of other species participating in mixed species foraging flocks (Freeberg et al. 2017) and predator mobs (Sieving et al. in press). In both cases, social brain evolution creates clear pathways to both high cognitive and vocal complexity demonstrated by TUTI and other parids (Langham et al. 2006; Urhan et al. 2026) – well known community informants (Carlson et al. 2020). Indeed, parids as a group have wide-ranging impacts on population and community-level behavior (Forsman et al. 1998; Jones & Sieving 2019) and even winter survival of associated species (Dolby & Grubb 1998).

Here we show also that for information from competent informants to manifest in appropriate community responses, competent eavesdropper species, including “non-social” species like cardinals are necessary. In the simplest sense, if eavesdroppers respond appropriately to the signals of community informants, this (by definition) defines the informant and demonstrates competent eavesdropping (Carlson et al. 2020). Certainly, there should be strong natural selection for any individual prey to learn the anti-predator calls of their local community informant(s) through social learning from heterospecifics throughout life, parents during even short post-fledging parental care periods (Seppänen & Forsman 2007; Farine et al. 2015; Carlson et al. 2019; Lind et al. 2019). Some essential anti-predator behaviors in birds develop naturally from a few days after fledging (basic avoidance; Tome & Denac 2012) and some parental guidance occurs (Griesser et al. 2016; Dutour et al. 2019), but social learning from the heterospecific community during a bird’s lifetime appears to be the most viable explanation for the training of competent eavesdroppers (Szymkowiak 2024). Indeed, most mobbing aggregations – where many species of scolding avian prey gather around a predator – are attracted by the mobbing calls of a community informant and these common aggregations likely serve as training grounds for community eavesdroppers (Curio et al. 1978). because the accurate discrimination of graded, intermediate-risk calls (GH Owl vs. Screech Owl) relies on sequential evaluation and sensory discrimination, these behaviors are likely learned and should strengthen with age and localized experience. In contrast, the unique, privileged status of the ZZ call in information processing leads to clear, testable predictions. Specifically, because the ZZ call bypasses complex cognitive evaluation, the attenuation of state-dependent behaviors under this threat should be highly robust and present even in naïve individuals or populations with consistent raptor exposure.

### Caveats and conclusion

We note the lack of individual identification and key measurements (e.g., of weight, personality types, etc), information which could help harness the individual level variation and whether the pre-playback feeding variable (PSE) could represent transient states (such as immediate hunger) or static individual behavioral types (e.g., proactive/reactive "personalities"). Also, since Blank and Contact trials contained zero fleeing events, the estimated state-dependence curves under those baseline assumptions are weakly constrained in the hazard model and must be interpreted with caution.

Despite these data constraints, our joint latent-risk framework—by linking multiple behaviors and accounting for excess non-feeding events—provides convergent, process-based evidence that cardinals actively exploit the associated information carried by heterospecific alarms to update their latent estimation of threat. This approach successfully captures a qualitative behavioral transition between the decoupled mental processing of sequential, graded threats—where correlational information serves as an ultimate structuring cause for adaptive vigilance—and the rapid, state-independent defenses triggered by highest-risk, instantaneous warning calls. Future work accounting for the interplay between ontogenetic cognitive flexibility (individual-level learning and decision-making) and phylogenetically canalized autonomic defenses (genetically hard-wired survival reflexes) will provide a richer, more unified understanding of how evolutionary forces shape the split-second decisions that define survival in information-rich, multi-species communities.

## Supporting information

Supplementary model and outputs

## ACKNOWLEDGEMENTS

Thank you to Colette St. Mary for her guidance with the dynamic optimization model. SH collected the field data as part of her undergraduate thesis; KS and SH designed the field study and drafted the initial manuscript; PC edited the initial draft and contributed insights during initial discussions and drafting stages; NM modeled the data and dynamic optimization to craft the central conceptual contribution of this paper and edited the initial draft.

