## Supplementary model and outputs for "Feed, Flee, Or Fact-Find: Modeling Latent Cognitive-mechanistic Responses to Heterospecific Alarm Calls for Foraging Cardinals"

### Dynamic Optimization

Trade-offs exist for all organisms capable of utilizing from a set of both fixed and plastic strategies. Among a finite set of possible strategies, we consider those which lead to an organism's greatest future fitness to be the "optimal strategy". Optimal strategies can vary from individual to individual as they are often highly dependent on an organism's physiological state as well as the state of an organism's environment. Likewise, physiological and environmental states often vary through time, and optimal strategies may vary as well. Dynamic optimization models use backward iteration from a terminal fitness function to determine which time-specific strategies maximize an organism's fitness given its current state or states. For a full description of these models and how they work, see Clark and Mangel (2000).

Our model considers two dynamic state variables, number of seeds eaten and perceived predation risk, such that fitness increases linearly with the number of seeds eaten and decreases proportionally to increasing risk. Time operates in one second units for a total of one minute, such that $T_{max} = 60$.

Mathematically, the terminal fitness function is,

$$\varphi(x) = 20x \quad . \tag{1}$$

where $x$ is the number of total seeds eaten and $p$ is the time-specific probability for risk of predation, as perceived by an individual.

We assume risk can never exceed 100% and therefore accumulates logistically through time such that,

$$p(t+1) = \frac{1}{\left(1 + \frac{1-p(t)}{p(t)} e^{-rt}\right)} \quad , \tag{2}$$

where $r$ is the rate of natural increase, and the parameter estimated for each call type. As $r$ approaches zero, accumulation of risk is approximately linear over the relevant time range.

At any given time, $t$, from 1 to 59, individuals can choose one of three strategies, **eat** which increases the number of total seeds eaten by one, **look** which decreases the perceived risk of predation ($R_p$) by 0.5, or **flee** which decreases risk of predation to 0.01. Thus, the set of possible strategies (S) is,

$$S = \{Eat, Look, Flee\}. \tag{3}$$

Each behavior carries a time cost which inhibits the ability to consume more seeds. For example, fleeing extends from time $t$ to the end of the time horizon, $T_{max}$, leaving no additional opportunity to consume seeds. Therefore, the number of seeds eaten at $T_{max}$ is equal to the number of seeds eaten when an individual flees. Likewise, looking up takes three seconds. Therefore, if an individual chooses to look up at time $t$, they cannot engage in any other strategy between time $t$

and $t + 3$. Finally, eating seeds takes time as well. Observations showed seed handling time to be six seconds on average. Thus an individual eating a seed at time $t$ cannot utilize other strategies between $t$ and $t + 6$.

Backward iteration of the dynamic optimization model operates according to the following,

Elements of the tensor/array **F** represents fitness through time and consists of three dimensions, time and the two state variables mentioned above, such that,

$$\boldsymbol{F}(x, p, T_{max}) = \varphi(x) \, , \tag{4}$$

where, again, $x$ represents the number of seeds eaten and $p$ represents perceived risk.

The set of functions $v_s$ represent expected future fitness for a given strategy $s$ belonging to the set $S$, for the same time range and set of state variables. That is,

$$v_s(x, p, t) = (1 - p) * \boldsymbol{F}(x_s, p_s, t_s) \;\; , \tag{5}$$

where $x_s$, $p_s$, and $t_s$ are the state variable changes for energy, risk, and time, respectively given strategy $s$.

Specifically,

$$v_{eat}(x, p, t) = (1 - p) * \boldsymbol{F}(x + 1, p(t + 1), t + 6) \, , \tag{6}$$

$$v_{look}(x, p, t) = (1 - p) * \; \boldsymbol{F}(x, 0.5p, t + 3) \, , \tag{7}$$

and,

$$v_{flee}(x, p, t) = \boldsymbol{F}(x, 0.001, T_{max}) \, . \tag{8}$$

The optimal strategy tensor, **D**, given the state of variables x and p, at time = $t$ is the strategy, $s$, that maximizes the function $v_s$. That is,

$$\boldsymbol{D}(x, p, t) = \max\big(v_s(x, p, t)\big) \;\; for \; all \; s \in S \;\; . \tag{9}$$

Optimal strategy tensors for different risk accumulation rates, $r$, where generated then used to determine strategies implemented in the forward simulations which are described below.

### Forward Simulation

We used forward simulations to evaluate behavioral outcomes under the optimal strategies generated by the dynamic optimization model. Initial perceived predation risk was fixed at 0.001, while initial energetic state (number of seeds previously consumed) varied among simulations. For each rate of risk accumulation, we calculated the corresponding optimal strategy tensor, ($\boldsymbol{D}_r$), and used this to determine individual behavioral decisions throughout the 60-second simulation.

At each time step, individuals selected one of three possible behaviors (eat, look, or flee) according to the optimal strategy associated with their current energetic state, perceived predation risk, and time. Individuals were temporarily prevented from making additional behavioral decisions while completing the time costs associated with eating or looking. Upon completion of an eating event, the individual's seed store increased by one with energetic state constrained to a maximum of 18 seeds, corresponding to the upper boundary of the dynamic optimization model. Likewise, perceived predation risk was updated according to equation 2. Following a looking event, perceived predation risk was reduced by 50% of its previous value, limited to 0.001. Individuals also accumulated one additional look for each looking event. In both cases, the resulting seed and risk state was used to determine subsequent behavioral decisions. Fleeing was treated as an absorbing state: individuals that fled were removed from subsequent behavioral iterations, and their time of departure was recorded. Individuals that did not flee during the 60-second simulation were assigned a departure time of 60 seconds.

We conducted simulations across the range of risk accumulation rates and initial energetic states. For each simulation, we recorded the total number of seeds consumed, total number of looking events, and time to flee. These outcomes were subsequently used to evaluate how initial energetic state and the rate of perceived risk accumulation influenced behavioral decisions and their resulting fitness consequences.

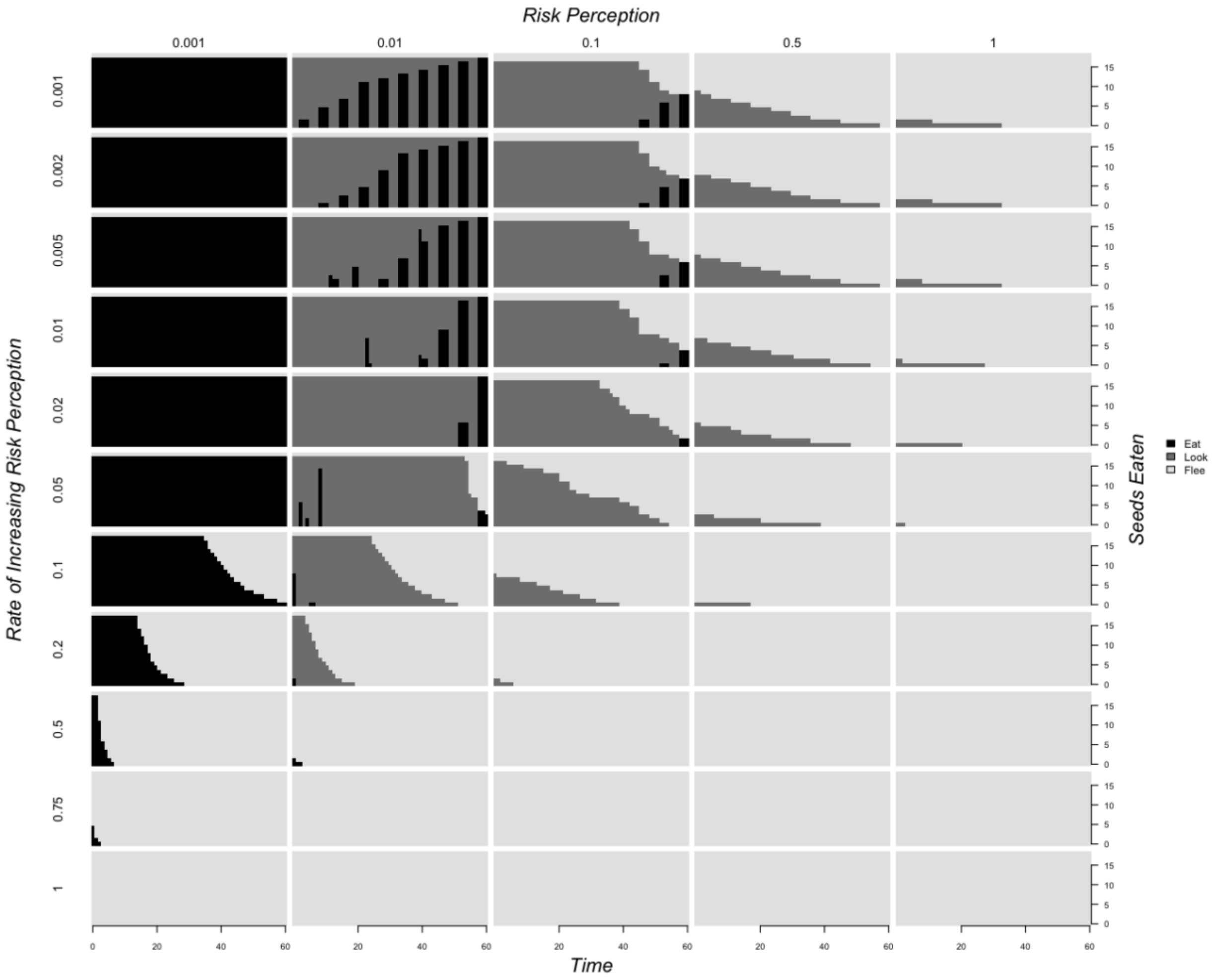


Supplementary Figure 1. Optimal behavioral strategies eat (black), look (dark gray), and flee (light gray) given number of seeds eaten and current risk perception, for different rates of increasing risk perception.

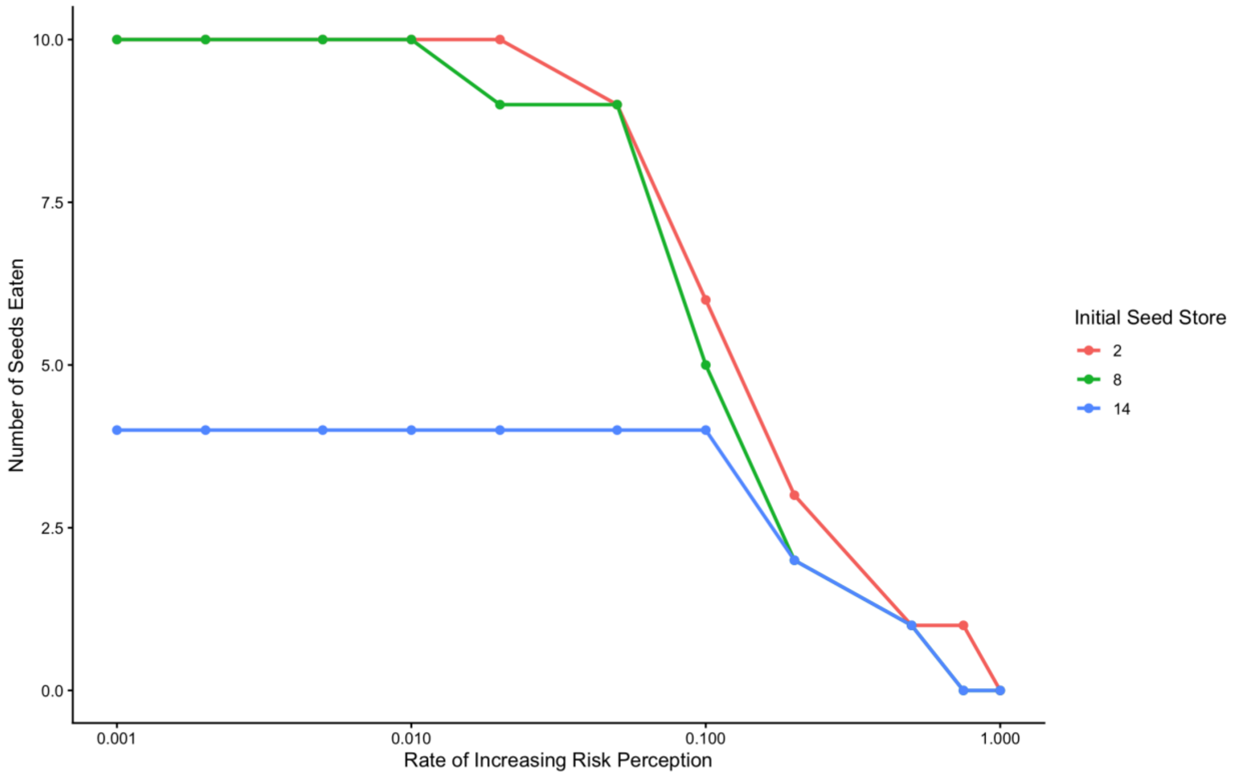


Supplementary Figure 2. Simulated number of seeds eaten for cardinals at $p(1) = 0.001$ against rate of increasing risk perception across three different initial seed stores low (2), medium (8), and (12).

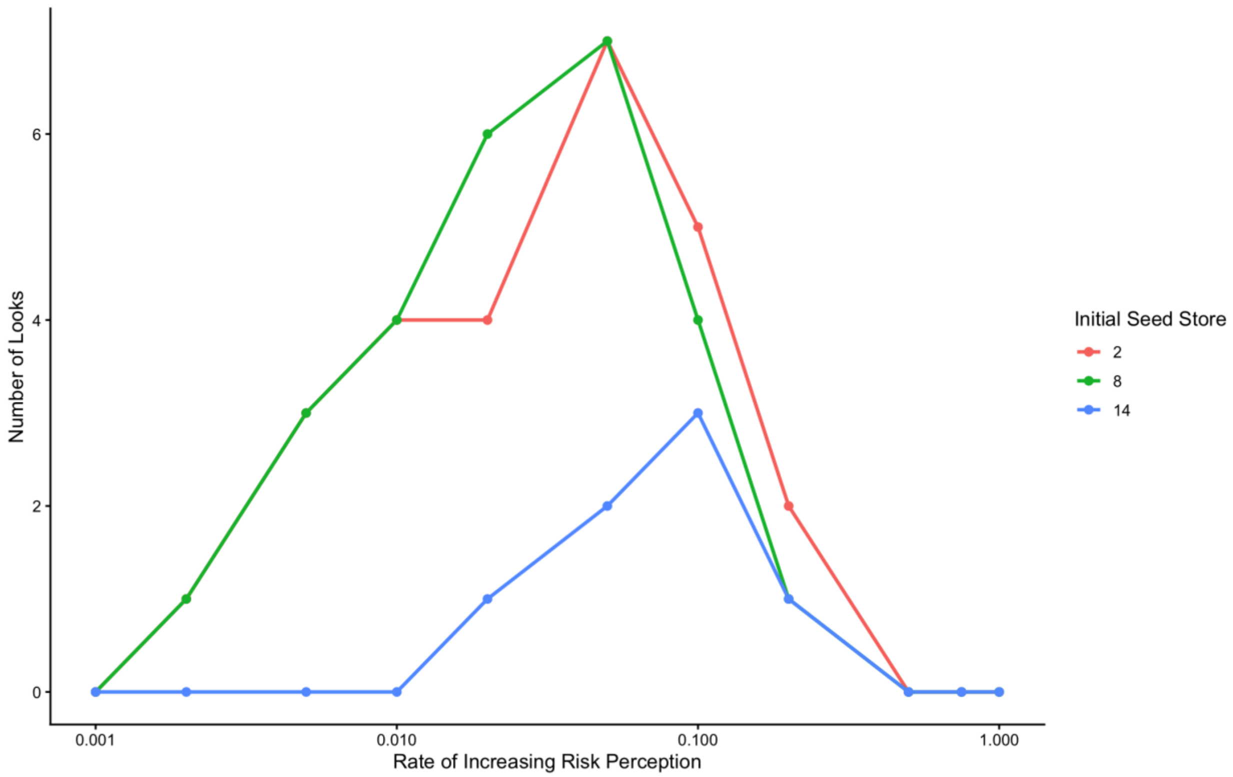


Supplementary Figure 3. Simulated number of seeds eaten for cardinals at $p(1) = 0.001$ against rate of increasing risk perception across different initial seed stores.

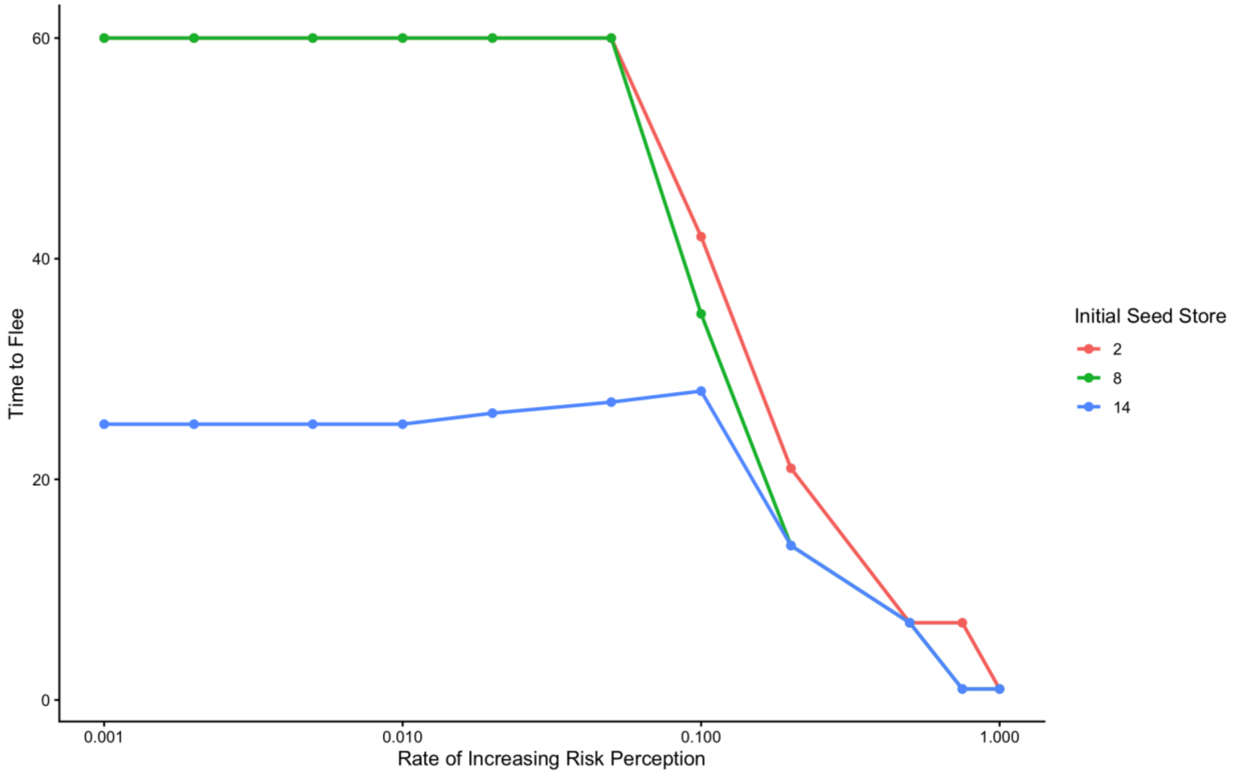


Supplementary Figure 4. Time to flee for cardinals at $p(1) = 0.001$ against rate of increasing risk perception across different initial seed stores.
